# Believer-Skeptic meets Actor-Critic: Rethinking the role basal ganglia pathways in decision-making and reinforcement learning

**DOI:** 10.1101/037085

**Authors:** Kyle Dunovan, Timothy Verstynen

## Abstract

The flexibility of behavioral control is a testament to the brain’s capacity for dynamically resolving uncertainty during goal-directed actions. This ability to select actions and learn from immediate feedback is driven by the dynamics of basal ganglia (BG) pathways. A growing body of empirical evidence conflicts with the traditional view that these pathways act as independent levers for facilitating (i.e., direct pathway) or suppressing (i.e., indirect pathway) motor output, suggesting instead that they engage in a dynamic competition during action decisions that computationally captures action uncertainty. Here we discuss the utility of encoding action uncertainty as a dynamic competition between opposing control pathways and provide evidence that this simple mechanism may have powerful implications for bridging neurocomputational theories of decision making and reinforcement learning.

## 1. Introduction

Consider the scenario of being presented with a plate of cookies. You first grapple with the decision as to whether or not you even want a cookie, depending on your fortitude at maintaining dietary goals. After a brief deliberation you decide to reach for a particularly delicious looking wafer, but along the way you notice that what you thought was a chocolate chip is in fact a spider resting on top, prompting you to reactively cancel your movement. The experience of seeing the spider will impact the certainty that you will reach for a cookie in the near future. This adaptability of both proactive (i.e., breaking your diet) and reactive (i.e., responding to the spider) behavioral control, in the face of multiple sources of uncertainty, is one of the most evolutionarily important functions of the mammalian brain.

Several lines of evidence point to a central role of cortical and basal ganglia (BG) circuits in modifying action decisions in dynamic environments; however, the mechanisms by which cortico-BG pathways encode uncertainty and adapt with experience remains controversial, fueled by a history of often inconsistent and sometimes paradoxical experimental findings. Central to this debate is the canonical model of the BG (Albin, Young, Penney, Roger, & Young, 1989; DeLong, 1990), where action selection is determined by the dynamics of three separate control pathways (Figure 1A): the direct pathway (the “Facilitation” signal) that facilitates motor output, the indirect pathway (“Suppression” signal) that suppresses motor output, and the hyper-direct pathway (“Braking” signal) that acts as a fast cancellation of facilitated motor decisions. According to the canonical model, all three pathways act as independent decision processes that regulate subsequent thalamic output to cortex (DeLong, 1990).

**Figure 1.**
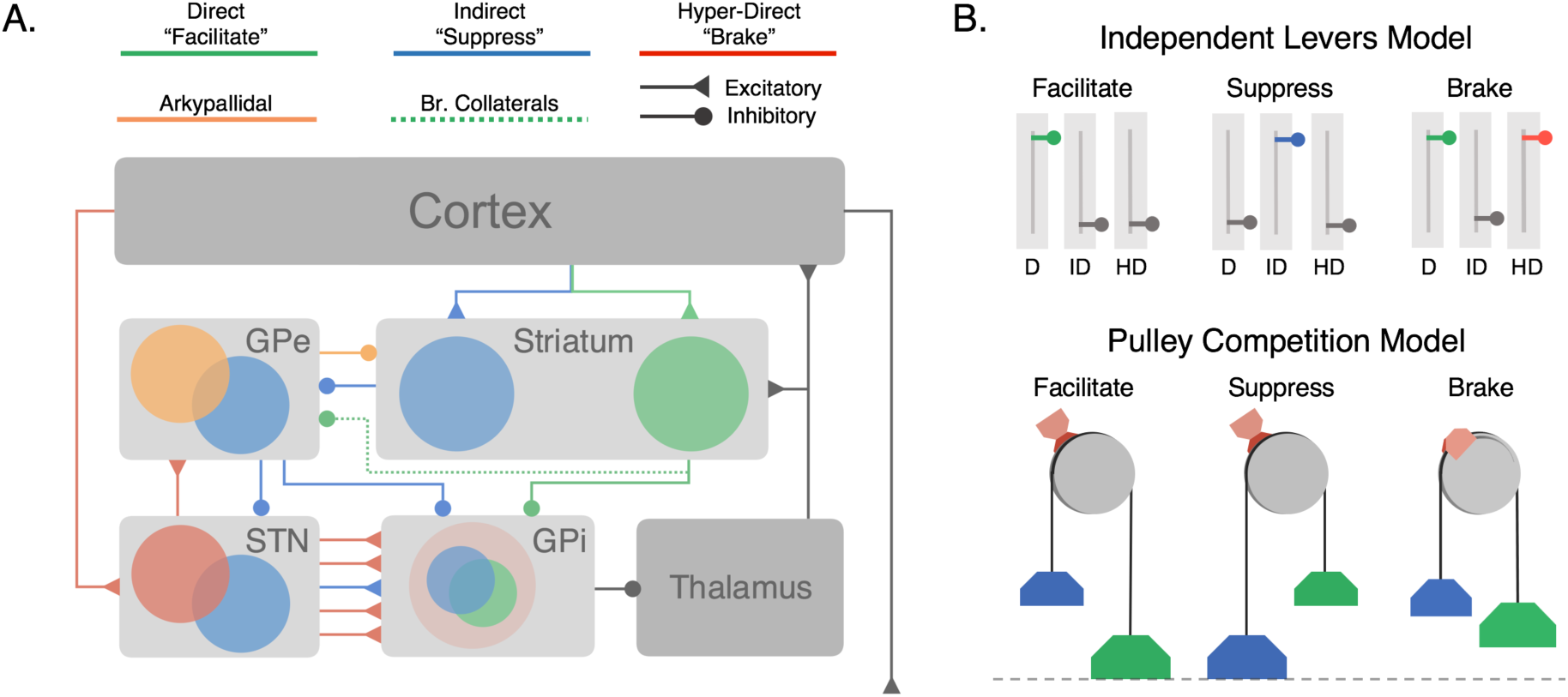
Architecture of cortico-BG pathways and hypothesized functional models. (A) Cortico-BG pathways including three major inputs to the striatal direct (green), indirect (blue) pathways, and the subthalamic hyper-direct (red) pathway. Bridging collaterals (green, dotted) connect the direct pathway to the indirect pathway via projections to the GPe. The arkypallidal pathway (orange) provides inhibitory feedback projections from the GPe to the striatum. Both the direct pathway (cortex-striatum-GPi) and “short” indirect pathway (cortex-striatum-GPe-GPi) form focused projections throughout the network corresponding to individual action channels. The “long” indirect pathway (cortex-striatum-GPe-STN-GPi) and hyper-direct pathway (cortex-STN-GPi) deliver diffuse excitatory inputs to the output nucleus. (**B**) Independent Levers Model (i.e., the canonical model) assumes that the direct (left, green), indirect (middle, blue), and hyperdirect (right, red) pathways are structurally and functionally segregated. Each pathway is operated in isolation for facilitating, suppressing, or braking motor output in the BG. (**C**) Pulley Model (i.e., Believer-Skeptic) assumes that the direct and indirect pathways compete throughout the BG (see Section 1), with the strength of each pathway acting as weights on opposing sides of a pulley. As activation in the direct pathway overpowers that of the indirect pathway, this imbalance accelerates the network toward “facilitation”, resulting in an executed action when the difference reaches a critical threshold (dotted line). In the event of a stop cue, the action can be reactively cancelled if the pulley brake (red brake pad) is activated before the direct-indirect difference reaches a critical threshold. The accelerating (e.g., nonlinear) dynamics of an imbalanced pulley lead to less efficacious braking when the network is pulled further toward action execution (e.g., longer brake streaks on pulley wheel). This dependency illustrates how proactive modulation of the direct-indirect balance may influence reactive stopping via activation of the hyper-direct pathway.

The architecture of the BG is such that each control pathway converges on a common output nucleus (the internal globus pallidus in humans), suggesting that at some level these pathways may interact. Indeed recent electrophysiological (Cazorla et al., 2014; Cui et al., 2013; Kress et al., 2013; Mallet et al., 2012), neuroimaging (Chikazoe et al., 2009; Jahfari et al., 2011, 2012), computational (Bahuguna, Aertsen, & Kumar, 2015; Dunovan, Lynch, Molesworth, & Verstynen, 2015; Gurney, Humphries, & Redgrave, 2015; Wei, Rubin, & Wang, 2015), and behavioral (Verbruggen, Stevens, & Chambers, 2014) findings have cast doubt on the traditional independent process framework, in favor of a dependent process model where all three pathways compete for basal ganglia output. These observations allude to a novel reconceptualization of basal ganglia pathways where the dynamics between all three pathways reflects a combination of "cooperative" and "competitive" states as a weighted function of learning and decision variables (Bahuguna et al., 2015; Cazorla et al., 2014; Dunovan et al., 2015; Gurney et al., 2015; Wei et al., 2015). This provides a theoretically valuable premise for characterizing BG involvement when adapting actions in uncertain environments (Wessel & Aron, 2013).

Here we explore the computational utility of a dependent process model of BG pathways. This review is partitioned into three sections. First, we provide an in depth summary of current debates regarding the role of the BG in inhibitory control. Next, we discuss recent advances relating computational models of decision-making and reinforcement learning to activity in cortico-BG networks. Finally, we propose a framework for synthesizing control, decision making, and learning within BG circuits, arguing that these pathways are best characterized by their ability to integrate uncertainty into goal-directed actions.

### 1.1 IntInteractions between direct and indirect pathways

.According to the canonical BG model, in the cookie scenario describe above the decision to reach for the cookie is driven by cortical activation of the direct pathway, whereas the decision to abstain is driven by activation of the indirect pathway. These two control signals are traditionally thought to occur in isolation of one another, such that upstream cortical regions either facilitate actions by activating the direct pathway or suppress actions by activating the indirect pathway (Hikida, Kimura, Wada, Funabiki, & Nakanishi, 2010). More recently, the canonical model has been revised to include a third ‘hyper-direct’ pathway in which cortical excitation of the subthalamic nucleus (STN) applies strong, diffuse suppression of action-facilitating signals in the direct pathway when a cue to stop (e.g., spider) is detected in the environment. This pathway is thought to race against action facilitating signals in the direct pathway in order to cancel an inappropriate or unnecessary action (Aron & Poldrack, 2006). Together the dynamics of the direct, indirect, and hyper-direct pathways form the basic building blocks of behavioral control in the canonical BG model.

The canonical BG model fundamentally assumes that all three control pathways run in parallel to each other and do not interact. Thus, the direct and indirect pathways may be viewed as two independent levers that are recruited in order to select appropriate actions that are in line with current behavioral goals (Figure 1B). This is often referred to as ‘proactive’ control (Braver, 2012). The hyper-direct pathway also acts as an independent lever, but one that is recruited ‘reactively’ upon detection of an environmental stop cue rather than endogenous goals (Aron & Poldrack, 2006). That is, the hyper-direct pathway acts as a safety brake for situations that require late action cancellation whereas the indirect pathway serves to selectively suppress actions which conflict with the current goals.

The notion that cortico-BG pathways operate as independent control mechanisms during action selection is reinforced by a large body of evidence demonstrating their opposing effects on motor output (see Albin et al., 1989 for review of basal ganglia motor circuitry and Calabresi, Picconi, Tozzi, Ghiglieri, & Di Filippo, 2014 for an updated view). Recently, Kravitz et al., (2010) showed that optogenetic stimulation of direct pathway medium spiny neurons (dMSN) facilitated locomotor behavior in mice, whereas stimulation of indirect pathway MSN’s (iMSN) led to motoric freezing. This was interpreted as strong evidence for the existence of structurally and functionally separate pathways for facilitating and suppressing movement. In contrast with the findings of Kravitz and colleagues (2010) a recent study by Cui et al., (2013) showed that both direct and indirect MSNs in the mouse dorsal striatum increase their firing just before contraversive movements. These findings provide the first clear evidence of a long theorized (Alexander & Crutcher, 1990; Mink, 1996), but empirically unfounded, action selection mechanism in the BG whereby cortical projections activate the direct pathway of a target action while simultaneously activating the indirect pathway of competing actions. Intuitively, this form of “center-surround” selection (Mink, 1996) becomes increasingly advantageous when there are many alternative actions from which to choose, acting as a safeguard against co-expression of multiple, interfering outputs. In this context, the observation that direct and indirect pathways are activated in unison marks an important discovery, but one that is not inconsistent with the independent levers model. Both pathways retain the same opposing influence over motor output and are operated independently and exclusively for each given action channel. Still these findings demonstrate that the direct and indirect pathways can be recruited in a complementary way via activation of the direct pathway for a single action (i.e. “center”) while activating the fashion indirect pathway of competing actions (i.e. “surround”).

This evidence of co-activation of direct and indirect pathways during action planning suggests that, rather than behaving as independent levers, the BG pathways may instead act like a pulley system, in which controlled actions arise from the competitive dynamics between direct and indirect pathways. In this novel conceptualization of BG pathways, the activation within the direct and indirect pathways can be represented by two weights, one on each side of the pulley (Figure 1C). As cortical inputs add to the weight of the direct pathway the pulley becomes increasingly imbalanced until a critical threshold is reached and an action can be executed. We should emphasize that, although the hyper-direct pathway may be viewed as a safety brake on the pulley that, if applied soon enough, it can prevent the weight of the direct pathway from overcoming the counterbalance of the indirect pathway weight and executing an action.

Architecturally there is ample evidence to suggest that BG pathways interact with each other. Most notably, all three pathways converge at the output nucleus of the BG, the internal segment of the globus pallidus (GPi) in humans and substantia nigra reticulata (SNr) in rodents. This region is generally considered to represent the locus of determination for action decisions. At rest the GPi tonically inhibits the thalamus, marking an important property of BG circuitry in that the default state of the network is suppressive. Thus, in order to elicit a motor output, the direct pathway must sufficiently suppress targeted cells in the GPi in order to release their inhibition on corresponding motor circuits (Mink, 1996). In situations requiring the inhibition of an action, indirect and hyperdirect pathways must strengthen pallido-thalamic inhibition to sufficiently override the effects of the direct pathway signals. Given that all three of the major BG pathways show signs of convergence in the GPi (Mathai & Smith, 2011; Smith, Bevan, Shink, & Bolam, 1998), it is easy to see how they could compete for a final output decision from the BG to the motor thalamus.

Recently another BG pathway was discovered that originates in a distinct population of cells within the external segment of the globus pallidus (GPe) and sends inhibitory feedback projections to the striatum (Mallet et al., 2012). These pallido-striatal feedback projections (i.e., the “arkypallidal” pathway, Figure 1A) preferentially synapse onto fast-spiking interneurons (FSIs) in the striatum rather than directly synapsing onto either of the major MSN subtypes. Given that FSIs form stronger connections with dMSN than iMSN populations (Mastro, Bouchard, Holt, & Gittis, 2014), the arkypallidal pathway may selectively facilitate BG output by reducing local inhibitory effects on dMSN output (Bahuguna et al., 2015). In addition to structural overlap between pathways within the basal ganglia, evidence now suggests that cortical projections to these pathways are not as segregated as previously thought, with both dMSN and iMSN subtypes receiving convergent thalamic (Huerta-Ocampo, Mena-Segovia, & Bolam, 2013) and cortical inputs (S. N. Haber, 2014; Kress et al., 2013; Wall, DeLaParra, Callaway, & Kreitzer, 2013). Still, there is a reliable tendency for prefrontal and frontal motor cortices to innervate the iMSNs and for sensory and limbic cortices to innervate dMSNs (Wall et al., 2013).

Finally, there is one architectural feature of BG circuits that has explicitly been shown to mediate an interaction between direct and indirect pathways: bridging collaterals are projections from the striatum that allow for D1 striatal cells, i.e., dMSNs, to project to the GPe and act as indirect pathway efferents (Y. Wu, Richard, & Parent, 2000). A recent study by Cazorla and colleagues (2014) showed that dMSN bridging collaterals terminating in the GPe are proliferated by promoting indirect pathway activity via D2R-upregulation in iMSN’s. The authors demonstrate through a series of experiments how experience-dependent changes in bridging collateral density alter the physiological and behavioral dynamics associated with direct and indirect pathway activation. In stark contrast with independent levers model, Cazorla et al., (2014) found that optogenetic stimulation of the direct pathway coincided with a moderate number of inhibited cells in the GPe in control mice, demonstrating clear interaction between direct and indirect pathways in normally developed animals. Remarkably, this effect became more salient with the activity-dependent proliferation of collaterals and actually reversed the effect of the direct pathway activation on behavior – suppressing locomotion rather than facilitating it.

One major implication of the Cazorla et al. (2014) study is that frequently suppressed actions, such as those that are costly or uncertain, become more difficult to execute as cortical activation of the direct pathway is restricted by proliferated dMSN collaterals into the indirect pathway. This functional link between direct and indirect pathways could potentially explain numerous conflicting findings in electrophysiological and human neuroimaging studies. For instance, both pathways, when stimulated in isolation, lead to heterogeneous (increased and decreased) changes in the firing of downstream GPi/SNr cells (Freeze, Kravitz, Hammack, Berke, & Kreitzer, 2013), whereas others (Kravitz et al., 2010)have demonstrated clearly opposing behavioral effects following direct (e.g., facilitation) and indirect (e.g., suppression) pathway stimulation. These seemingly inconsistent findings can be reconciled by revising the canonical model to incorporate cross-talk between the direct and indirect pathways, either through direct-pathway bridging collaterals or through arkypallidal feedback projections to the striatum. Finally, human neuroimaging studies of response inhibition have concluded that proactive control is singularly driven by cortical activation of striatal indirect pathway (Majid, Cai, Corey-Bloom, & Aron, 2013). The findings by Cazorla et al. (2014), in addition to many of the findings discussed above, strongly caution against the notion that proactive control arises from exclusive engagement of the indirect pathway or that modulation of this control is limited to cortical sources.

## 2. Believer-Skeptic: Encoding Uncertainty as a Dynamic Competition

The studies discussed thus far provide evidence against the independent lever model of cortico-BG pathways and instead favor a model in which these pathways engage in a dynamic competition: as activity increases in one of the pathways the balance is upset and the network accelerates toward motor-facilitating or motor-suppressing state. Seen in this light, this direct-indirect competition represents a potentially important decision-making mechanism whereby multiple sources of uncertainty can be weighed and integrated before choosing between potential actions. In this way, the direct-indirect competition implements a decision by weighing the arguments of a Believer (e.g., direct pathway) against those of a Skeptic (e.g., indirect pathway). Because, the default state of the BG is heavily motor suppressing (Bahuguna et al., 2015), the burden of proof is on the Believer and thus, actions are only executed when the accrued evidence sufficiently reduces the Skeptic's uncertainty. Here, we show that the competition between the direct and indirect pathways can be formalized by the dynamics of a simplified neural network model of cortico-BG pathways and mapped onto parameters of accumulator models of decision-making. From this, we argue that the competitive nature of cortico-BG pathways is a critical feature for encoding uncertainty and adapting behavior in changing environments.

### 2.1 Competing BG pathways encode decision uncertainty

Computational models of decision-making predominantly fall within the broader class of accumulation-to-bound models, in which a decision is computed by accumulating the evidence for one choice over another until a threshold is met and a choice can be made. When deciding between two alternative hypotheses, or choices, the optimal rate of accumulation and criterion setting for maximizing speed and accuracy is described by the Drift-Diffusion Model (DDM) (Ratcliff, 1978; see Ratcliff & McKoon, 2008 for a review). Successful application of this model to a broad spectrum of behavioral phenomena has established the DDM as the archetypal model of decision-making, providing a parsimonious framework capable of describing both the speed and accuracy of decision behavior. By fitting models to behavioral data, response-time and accuracy measures are decomposed into hypothesized subcomponents of their generative mechanism, quantified by the model’s parameters. Parameters can then be used to extract neural activity related to individual subcomponents of the decision mechanism. While significant progress has been made by leveraging stochastic accumulator models to aid in the prediction and interpretation of data in experimental neuroscience, it remains an open question at what level of neural processing (i.e., single neurons, local circuits, networks) these parameters are realized in the brain.

Studies investigating the neural basis of decision-making have largely focused on frontal and parietal systems, following from early observations that single-neurons in these regions appear to display the same ramp-to-threshold characteristics as the DDM. More recently, it has become clear that the neural processes involved in decision-making are much more distributed than previously thought, suggesting that decision variables are tracked by populations of neurons (Park, Meister, Huk, & Pillow, 2014) at both the cortical (Heitz & Schall, 2012, 2013) and subcortical (Ding & Gold, 2012b, 2013) levels. Indeed, mounting evidence points to the BG as a critical part of the decision network, serving as a convergence zone for contextual and sensory information prior to decision commitment (Ding & Gold, 2012b; Dunovan et al., 2015; Keuken et al., 2015; Nagano-Saito et al., 2012; Wei et al., 2015; Yanike & Ferrera, 2014). Most of the cortical regions that have been implicated in the evidence accumulation process send direct projections into the BG (Averbeck, Lehman, Jacobson, & Haber, 2014; Draganski et al., 2008; S. Haber, Kunishio, Mizobuchi, & Lynd-Balta, 1995; Jarbo & Verstynen, 2015; Verstynen, 2014), as do context‐ and performance-monitoring regions (Forstmann et al., 2012; S. Haber et al., 1995; Haynes & Haber, 2013; King et al., 2012). This convergence of cortically distributed decision signals into the BG adds credence to the growing body of evidence suggesting this network is critical for imposing a threshold on accumulating decision evidence (Bahuguna et al., 2015; Bogacz, Wagenmakers, Forstmann, & Nieuwenhuis, 2010; Cavanagh et al., 2011; Forstmann et al., 2008; Frank et al., 2015; Lo & Wang, 2006; Mansfield, Karayanidis, Jamadar, Heathcote, & Forstmann, 2011; Wei et al., 2015).

It is important to note that, in contrast with the DDM-like ramping of cortical accumulators, the neural implementation of a decision threshold is unlikely to present in such a straightforward manner (Heitz & Schall, 2013; Simen, 2012). Changes in the decision threshold of the DDM can capture decision-related computations that occur at various stages of processing; for instance, a decrease in the DDM threshold can describe the behavioral effects of indiscriminately increasing the baseline of evidence for both alternatives prior to sensory input. Furthermore, a shift in the baseline of evidence may reflect priming in lowlevel sensory regions, in downstream evidence accumulators, in cortical and subcortical motor circuits, or in some combination of these domains. Indeed, Heitz and Schall, (2012, 2013) have shown in a series of computational and electrophysiological studies that, behavioral adjustments optimally explained by a change in decision threshold in standard accumulator models arise from a combination of parameter changes in the of neurons in the frontal eye fields (FEF). According to these findings, the representation of the decision threshold in standard accumulator models is best thought of as an abstraction of more sophisticated network dynamics underlying speed-accuracy tradeoffs. Therefore, it is useful for the purposes of this review to clarify the meaning of *decision threshold* in the abstract sense so as to distinguish this meaning from the mechanism by which it is theoretically modeled or neurologically implemented. At a conceptual level, a decision threshold can be thought of as the “switch” or “latch” mechanism responsible for transitioning from an accumulation state to an action execution state. In contrast to the notion of a threshold as the ‘upper limit’ or ‘criterion boundary’ placed on evidence accumulation, switches are dynamic processes themselves and can be adjusted to be more or less sensitive to perturbation.

Converging electrophysiological (Ding & Gold, 2012a; Schall, Purcell, Heitz, Logan, & Palmeri, 2011) and computational (Simen, 2012; Standage, Blohm, & Dorris, 2014) evidence suggests that competing populations of neurons can implement a transition threshold in the presence of sufficient nonlinearity in the competitive inhibition between populations. For instance, Schall and colleagues (2011) proposed the gated-accumulator model to account for the cross-inhibition between target and distractor populations in the FEF. These authors trained macaques to maximize reward by emphasizing the speed or accuracy of their performance in a visual search task based on a prior cue. Behavioral speed-accuracy tradeoffs were well described by a traditional accumulator model allowing only the threshold to vary across conditions. However, recordings in choice-selective frontal eye field (FEF) neurons displayed simultaneous changes in the baseline, onset, and rate of firing as a function of decision policy. Consistent with this, Wei, Rubin, and Wang (2014) recently showed that competitive dynamics between the direct and indirect pathways in a spiking neural network of the BG could be tuned to strategically adjust the decision threshold. In their model, changing the synaptic efficacy of indirect pathway output from the striatum to the GPe effectively modulated the threshold at which accumulating cortico-striatal inputs produced an action. Thus, rather than manifesting as a change in the RT-locked firing rate of cortical accumulators (as might be expected if neural decision thresholds were implemented as in the DDM), this model showed that BG circuitry can approximate the same mechanism by modulating balance of the direct and indirect pathways.

Similar in concept to the gated accumulator model, consider the simple neural network shown in Figure 2A, composed of two competing neural populations with recurrent excitatory connections. The mutual inhibitory connections between populations of direct and indirect units, in combination with recurrent self-excitation, leads to a non-linear change in the separation of their firing rates over time. The point in time at which this separation occurs marks the ‘gate’ in the gated accumulator model. Network properties that promote early gating correspond to a lower threshold in the traditional DDM. That is, they both reduce allotted time for evidence to be gathered. On the other hand, the effective threshold can be “raised” by increasing the time constant of evidence accumulation, reflected in the network as a delayed gate or more gradual separation of competing population activity.

**Figure 2.**
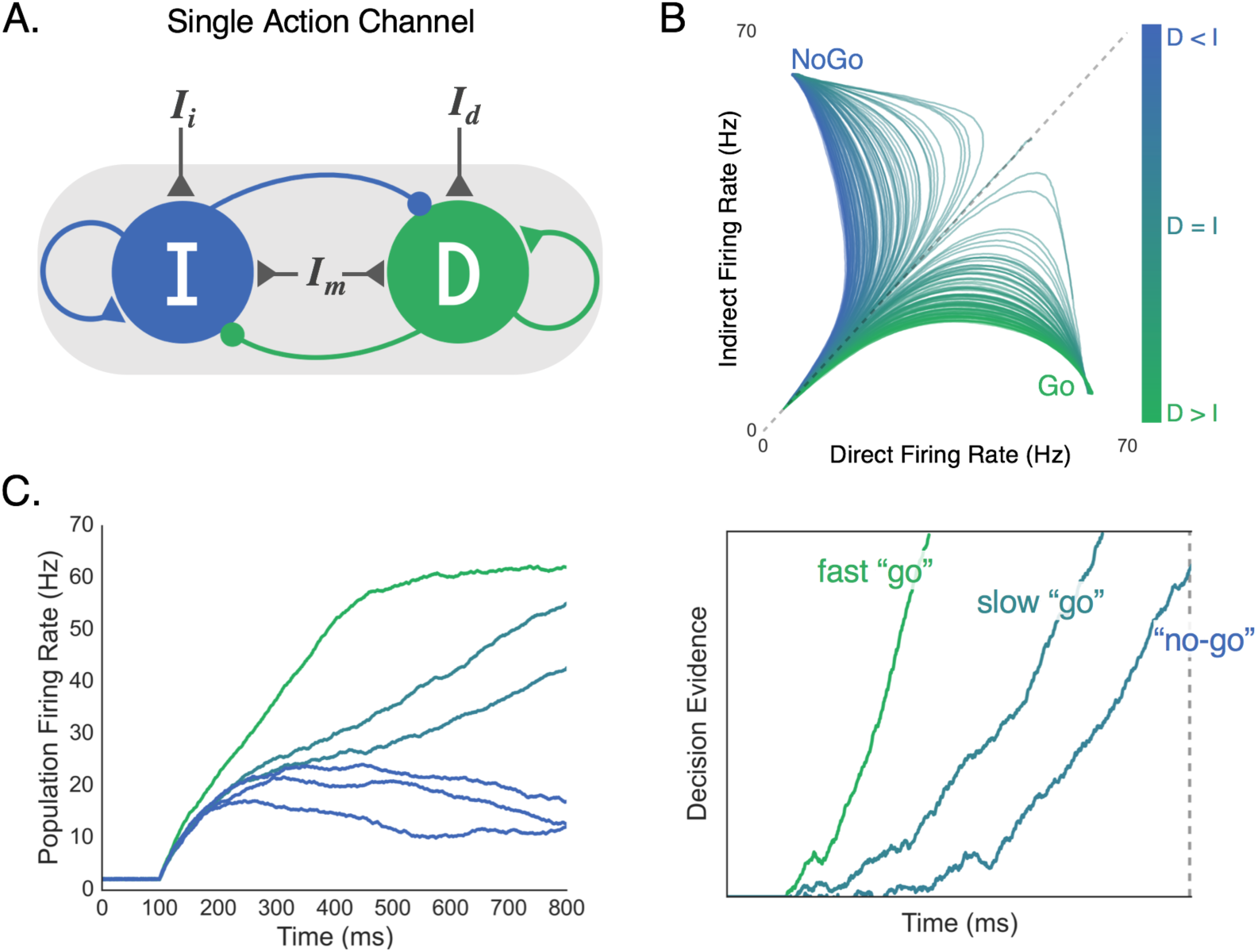
Believer-Skeptic framework and competition between the direct and indirect pathways in attractor and accumulator space. **(A)** The direct (D) and indirect (I) pathways are modeled as two competing (i.e., mutual inhibition) accumulators with recurrent self-excitation reflecting population attractor dynamics. Selective input to the direct (I_d_) and indirect (I_i_) pathways is weighted (not shown) and summed with input from a modulatory (non-selective) population (I_m_) which controls the baseline excitability of the network. **(B)**The firing rate of indirect (vertical axis) and direct (horizontal axis) pathways as a function of different ratios of pathway activation. Greater activation of the indirect pathway leads to fast attraction towards a NoGo state (blue, motor suppressing), whereas greater activation of the direct pathway attracts the network toward a Go state (green, motor facilitating). **(C)** Left panel: direct (higher firing rates) and indirect pathway (lower lower firing rates) firing rates plotted across time for different ratios of input (I_d_:I_i_). Right panel: accumulation of decision evidence toward an execution threshold, reflecting the normalized difference of the direct and indirect pathways in the left panel. High I_d_:I_i_ ratio accelerates the rate of evidence accumulation, leading to a fast “go” response. As this ratio is reduced, weaker attraction by the direct pathway (left, bluish-green) manifests as a slower rate of accumulation, producing a “no-go” decision when evidence fails to reach threshold by a deadline.

While the gated accumulator model was originally developed to capture activity in target and distractor populations of cortical neurons (Schall et al., 2011), we propose that a similar threshold mechanism is implemented by a competition between direct and indirect pathways. In this reduced form of the model, the respective strength of each pathway is determined by several factors, including the amount of cortical input to each population, the weight applied to those inputs (e.g., cortico-striatal synaptic efficacy), and the overall excitability of the network based on non-specific modulatory inputs. As mentioned above, others have postulated that co-activation of direct and indirect MSNs represents a center-surround action selection mechanism in which the direct pathway of a target action is activated along with the indirect pathway is activated for alternative actions (Cui et al., 2013; Friend & Kravitz, 2014). We propose that both pathways are activated for each individual action, but to varying degrees such that the ratio of direct-to-indirect activity is optimized during goal-directed learning. Thus, rather than the two populations in Figure 2A representing target and distractor stimuli, they represent a single action channel composed of a Believer population that competes with a Skeptic population.

The idea that the certainty of individual actions might be encoded by co-activation of direct and indirect pathways has been proposed by previous models of BG pathways (Brown, Bullock, & Grossberg, 2004; Schroll & Hamker, 2013; Wiecki & Frank, 2013); however, strong empirical evidence for this competition is limited to recent electrophysiological studies. This may be due, in part, to the fact that dMSN and iMSN’s are often not distinguished in single-unit studies of decision-related activity in striatum (Ding & Gold, 2010, 2012b, 2013). This is why the findings by Cui et al., (2014) have sparked excitement about the implications for sequential action selection facilitated by switching on the direct pathway for the target action (i.e. center) while activating the indirect pathway for a field of to-be selected actions (i.e. surround). However, a recent computational study also suggests that both pathways must be active to a controlled degree in order to optimize action selection in a goaldirected manner, otherwise no actions or too many actions are selected (Gurney et al., 2015). Under this assumption, a center-surround mechanism can still arise in which a target action enjoys a greater direct-to-indirect ratio than surrounding actions. In fact, there is good reason to think that actions are selected through a combination of center-surround suppression and the action-specific balance of facilitation and suppression. We elaborate more on this in the following section.

### 2.2 Linking Neural Competition to Accumulator Models

The Believer-Skeptic framework presented here proposes that cortico-BG pathways implement a decision threshold as a dynamic competition of action facilitating and suppressing network states. While we propose this to be a more neurally plausible mechanism of threshold implementation than that presented in the DDM, this is not to say that model abstraction is not useful. In fact, it is necessary for developing quantitative theories that can be meaningfully parameterized at cognitive and behavioral levels of description. In order for these models to be applied to neural data there must be an appreciation for the mapping between cognitive parameters and the more complex neural processes that they represent.

Within the standard DDM, ‘competition’ is inherently captured by the accumulating decision process where each step up or down represents the instantaneous evaluation of two competing hypotheses: an action decision and its null alternative. In the context of basic perceptual decisions, stimuli with high signal-to-noise ratio (SNR) produce faster rates of evidence accumulation toward a decision boundary, and are thus recognized faster and more reliably than noisy stimuli. This is an important point to emphasize, as the unidirectional change in the speed and accuracy of decisions is what fundamentally distinguishes a change in drift-rate from a change in the decision threshold in the standard DDM. As hinted at earlier the decision process can instead be reparameterized to reflect different hypotheses regarding the neural processes responsible for integrating contextual information with sensory evidence (Standage et al., 2014). In the Believer-Skeptic framework, contextual information and sensory evidence converge as weighted cortico-striatal inputs to the direct and indirect pathways of a single action channel (Fig. 2A). The strong recurrent dynamics within each pathway lead to bistability in the network output (Fig. 2B), an important property for implementing a switch between two states. Even when the weighted input to each pathway is comparable, small amounts of noise can disrupt the balance enough to cause a state transition given sufficient self-excitation. As a result, both pathways initially increase their firing rate then diverge as activation in one pathway supersedes and inhibits the other, switching the network toward a ‘Go’ or ‘NoGo’ attractor state (Fig. 2B). Thus, rather than the sensory driven drift-rate of the DDM, the moment-to-moment competition between alternative hypotheses in the Believer-Skeptic framework is driven by a weighted combination of contextual and sensory information. This form of competition can be seen in Figure 2C, in which Go-NoGo decisions are made by accumulating the output (right panel) of the direct-indirect competition (left panel) under different levels of contextual uncertainty. When action uncertainty is low, the network is accelerated toward a “Go” state (Figure 2B) by stronger activation of the direct pathway, causing a faster accumulation of decision evidence towards a fixed execution threshold. Neurophysiologically, the fixed threshold in this context can be conceptualized as the level of pallidal suppression required to sufficiently disinhibit the thalamus so that an action is executed.

We recently proposed a modified accumulator framework motivated by the general control dynamics of the Believer-Skeptic network in Figure 2, where action decisions are executed by accumulating evidence toward a fixed threshold in the presence of dynamic gain. In our so-called dependent process model, we found that contextual information (i.e. cued probability of reward) modulates the drift-rate of the execution process (as seen in the right panel of Figure 2C). As action uncertainty increases the drift-rate is suppressed, producing a ‘no-go’ decision when this suppression prevents the decision process from reaching the execution threshold by the trial deadline (Dunovan et al., 2015). Based on the apparent structural overlap of cortico-BG pathways in the BG output nucleus (shown as overlapping red, blue, and green fields in the GPi of Figure 1A), we hypothesized that contextual modulation of competition between direct (i.e., Go) and indirect (i.e., NoGo) pathways should also influence the efficacy of the hyper-direct (i.e., Stop) pathway during reactive action cancellation (Jahfari et al., 2011, 2012; Jahfari, Stinear, Claffey, Verbruggen, & Aron, 2010), Indeed, behavioral fits to RT and choice data in a reactive stop-signal task favored a model in which contextual suppression of the execution drift-rate improves the efficacy of a nested but separate action cancellation process. Collectively, these findings show how the contextual uncertainty associated with a future action is not only critical for making a goal-directed decision about executing that action, but also complements the ability to reactively cancel it based on environmental feedback.

This dependent process model also captured physiological responses of BG pathways. By integrating the execution process across the trial window, we were able to capture the duration and magnitude of accumulating activity leading up to a decision. Integrating the execution process in this way effectively collapses the decision process into a single measure, similar to how the blood oxygenlevel-dependent (BOLD) signal would filter the neural activity generated by attractor network in Figure 2A. Consistent with the behavioral fits, we found that contextual modulation of the drift-rate was able to capture the pattern of BOLD activity in the thalamus (the primary output target of the BG pathways) during ‘go’ and ‘no-go’ decisions across varying degrees of uncertainty. This finding is consistent with single-unit recordings of neurons in the macaque motor thalamus which show a similar RT-dependent ramp in firing rate prior to action execution (M. Tanaka, 2007; Masaki Tanaka & Kunimatsu, 2011).

One interpretation of this finding is that pre-action ramping in the thalamus is driven by the differential activation of upstream direct and indirect pathways and thus contextual modulation of this signal occurs by changing the weights of specific cortico-striatal connections or by altering background excitability in the striatum. The hypothesis that the striatum is where contextual information comes to bear on decision evidence is often contrasted with the hypothesis that this is accomplished by the thresholding function of the STN (Bogacz et al., 2010). That is, a change in the slope of thalamic firing rates could be due to decay in the hyper-direct activation of the STN, allowing pallidal suppression by the direct pathway to disinhibit the thalamus at a proportional rate. The distinction between striatal and STN control over decision threshold is a critical one (Bogacz et al., 2010), as these structures have very different input-output motifs that hint at disparate functional roles. The input-output organization of the striatum is thought to be channel-specific, propagating individual action-commands from cortex to corresponding units in the globus pallidus external (GPe; indirect) and internal (GPi; direct) segments. The STN, on the other hand, receives converging afferents from cortex and the GPe and delivers diffuse excitatory drive to the GPi, suggesting this structure modulates the decision threshold in a non-specific manner for all actions under consideration.

Recently, another hypothesis has been proposed for the role of the STN in decision-making that both complements the role of the striatum in the Believer-Skeptic framework and distinguishes the functional relevance of indirect and hyper-direct activation of the STN. Bogacz & Gurney, (2007) presented a neural network model in which the STN normalizes activity in the GPi to accommodate different set sizes of alternative choices. In their model, sensory evidence for each alternative is fed into a corresponding action channel in the striatum in parallel with projections that activate the STN. As a result, the cortico-striatal activation within each individual channel of the GPi (i.e., representing candidate actions ‘A’, ‘B’, and ‘C’, for instance) is represented as a proportion of the evidence for each action relative to the total evidence for all actions under consideration. This model describes the general increase in RT associated with increasing the number of choices to be considered, indicative of a global increase in the threshold for all possible outcomes (Keuken et al., 2015). Another group found that removal of the STN from the network had similar effects on choice RTs as STN deep brain stimulation in treated Parkinson’s patients - selectively eliminating the delay in RT for low-probability stimuli (Antoniades et al., 2014).

The proposed thresholding and normalization functions of the STN are complementary with the Believer-Skeptic framework and can be dissociated from the hitherto-proposed role of the direct and indirect competition as a mechanism for encoding action uncertainty. The normalizing effect of STN output on pallidal inhibition emerges naturally under the assumption that all actions simultaneously engage both the direct and indirect pathways. That is, individual action uncertainty is encoded by the “short” indirect pathway from striatum to GPe to channel-specific populations in the GPi (see Figure 1; Schroll & Hamker, 2013) where the indirect pathway converges with action facilitating signals of the direct pathway. On the contrary, activation of the “long” indirect pathway, splitting off from GPe to the STN, leads to widespread excitatory increase in GPi firing. Under the assumption that both direct and indirect pathways are active for each action being considered, the net activation through the “long” indirect pathway has a normalizing effect on the basal GPi state, accommodating varied set sizes of alternative actions. Moreover, the relative uncertainty between actions is preserved regardless of hyper-direct perturbation of STN in the event of conflict detection. Increased hyperdirect activation of the STN would sacrifice the optimality of the normalizing constant it delivers to GPi, but only when that optimality is challenged by unanticipated conflict.

While the long-indirect and hyper-direct pathways likely play an important role in action selection, the within-channel competition of the direct and (short) indirect pathways is ultimately what determines which action is selected. For instance, in the context of a forced-choice perceptual decision, the transition between accumulation and execution is determined by the relative activation of two alternative action channels, each driven by separate pair of competing direct and indirect populations. This process is shown in Figure 3, where an observer must decide whether a noisy field of moving dots contains greater coherent leftward or rightward motion. Critically, a cue is displayed prior to each choice informing the observer which outcome is more likely to be correct on the upcoming trial. Previous work has shown that this predictive information is encoded by a concurrent increase in the baseline activity in the striatum, contralateral to the expected action, and modulatory regions of cortex, such as orbitofrontal cortex (OFC) and pre-supplementary motor area (preSMA) (Forstmann, Brown, Dutilh, Neumann, & Wagenmakers, 2010). When the cued probability is valid (i.e., correctly predicts the subsequent stimulus; Fig. 3A) the increase in baseline activity of the corresponding action channel causes the network to become more unstable, leading to faster gating upon descending input from cortical accumulators. However, when the cue is misleading, or invalid (Fig. 3B), this destabilization in the cued action action channel can lead to an incorrect response despite weak sensory evidence in favor of that choice. This speed-accuracy tradeoff is a widespread phenomenon that pervades all forms of decision-making. While numerous studies have found that functional and structural connectivity between preSMA and the striatum predicts with individual differences in the speed-accuracy tradeoff (Forstmann et al., 2010; Keuken, Langner, Eickhoff, Forstmann, & Neumann, 2014; van Maanen et al., 2011), the underlying mechanism by which modulatory cortical inputs influence action selection in the BG has remained unclear. The example here proposes one such mechanism and highlights an important prediction of the Believer-Skeptic framework, in which uncertainty associated with individual actions is encoded by the competition between corresponding direct and indirect pathways. Of course, this prediction will need to be more rigorously tested, both experimentally and through the use of more sophisticated computational models of BG circuitry.

**Figure 3.**
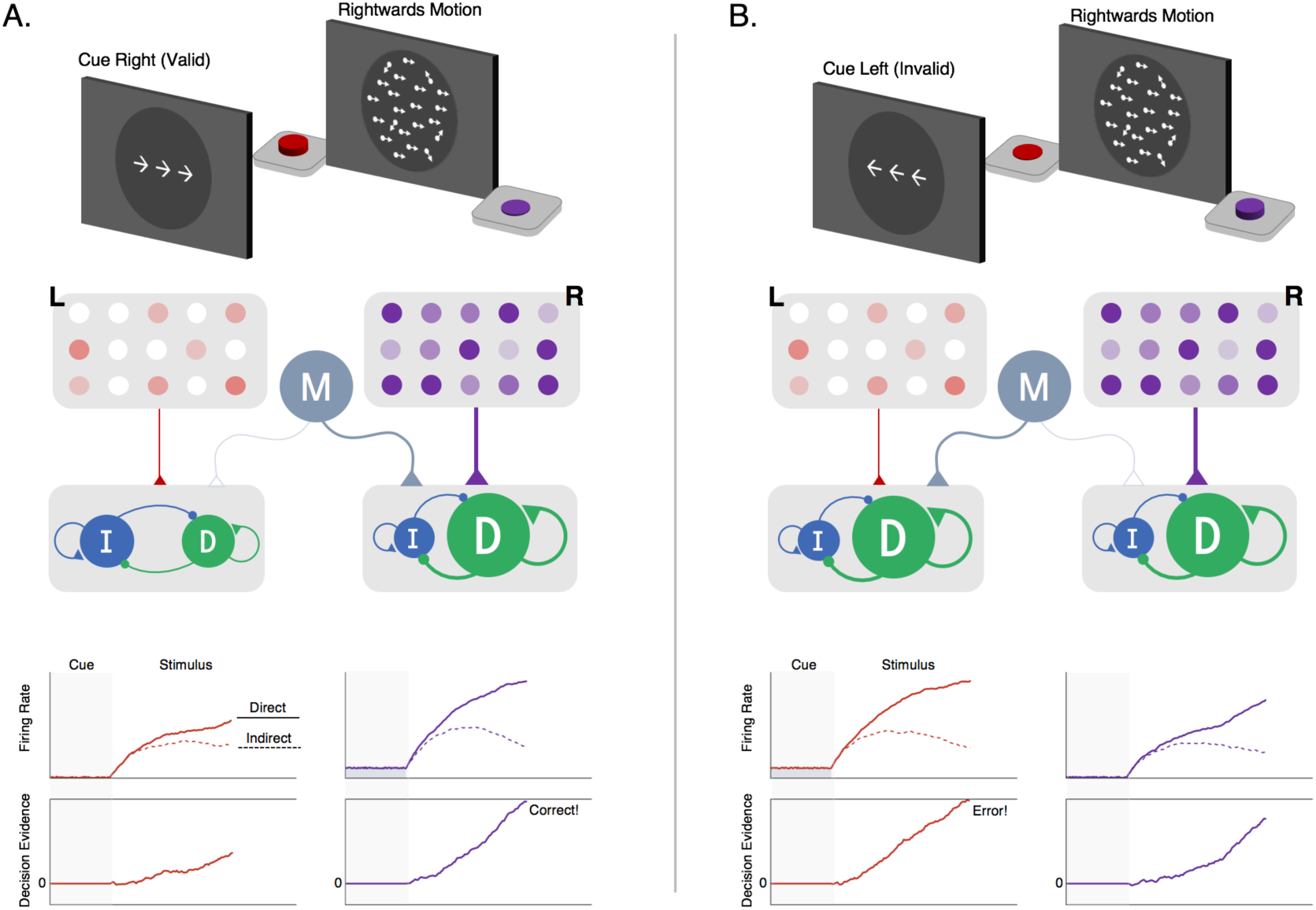
Within-channel competition and top-down modulation of expectations during perceptual decision-making. **(A)** Top: schematic of cue and stimulus epochs of random dot motion task on trial with valid predictive cue-stimulus combination. Middle: schematic of decision network. Left (L, red) and right (R, purple) motion-selective sensory populations gradually increase activity at a rate proportional to the strength of coherent motion in their preferred direction. Each sensory population sends excitatory input to a corresponding pair of direct and indirect populations representing left‐ and right-hand actions for reporting leftward and rightward motion decisions, respectively. Sensory inputs activate both pathways but with a bias favoring the direct pathway, reflecting the tendency for sensory inputs to the striatum to form more connections with dMSNs than iMSNs (Wall et al., 2013). A modulatory population (M, gray) delivers nonselective excitatory input to the pair of direct and indirect pathways encoding the anticipated action (i.e., action corresponding to the cue-predicted motion direction). Bottom-upper: firing rates of the direct (solid line) and indirect (dotted lines) populations for left‐ and right-hand actions. Bottom-lower: accumulation of the difference between direct and indirect firing-rates toward an execution threshold. The effect of cued expectations can be seen as an upwards shift in the baseline firing rates of the right-hand direct-indirect network, reflecting anticipatory input from the modulatory population. This increases the excitability of the network, causing a faster separation in the direct-indirect competition and a faster rise-to-threshold in the rightwards than leftwards decision variable, producing a correct response. **(B)** Top: same task as in the left panel but on a trial with an invalid predictive cue-stimulus combination. Middle: same decision network as in the left panel but with modulatory input delivered to the left-hand direct-indirect network as a result of cued expectations of a leftwards motion stimulus. Bottom: same layout as in the left panel. The invalid expectation signal destabilizes the direct-indirect competition, leading to a faster rise-to-threshold of the lefthand decision variable and an incorrect response.

In sum, the Believer-Skeptic framework provides a compelling account for the role of the BG in decision-making, demonstrating the computational utility for encoding action uncertainty in the competition between the direct and indirect pathways. This framework also provides a straightforward interpretation of the different roles of striatal and subthalamic modulation of the decision process. Non-specific background inputs to the striatum can adjust the speed-accuracy tradeoff in favor of quicker decision-making by promoting faster state attraction in response to input from sensory accumulators. Cortico-striatal mechanisms may also modulate the decision in outcome-specific ways by altering the balance of channel-specific activity in the direct and indirect pathways. This interpretation is consistent with human neuroimaging studies linking cortico-striatal activity to the facilitation of one choice at the expense of choosing another; for instance, by selectively increasing of the drift-rate or baseline evidence for an expected outcome (Dunovan et al., 2015; Forstmann et al., 2010). On the other hand, indirect pathway activation of the STN provides a normalizing constant to BG output by aggregating the activation of multiple action channels into diffuse projections to the GPi, whereas hyper-direct activation of the STN modulates the decision indiscriminately, buying time in the interest of accuracy (Forstmann et al., 2012; Frank et al., 2015). In the following and final section, we elaborate on how Believer-Skeptic dynamics of decisionmaking are complemented by the well-established role of the cortico-striatal circuits in mediating Actor-Critic reinforcement learning (RL).

## 3. Believer-Skeptic meets Actor-Critic

The idea that direct and indirect pathway competition may be a mechanism for encoding action uncertainty has profound implications not only for decision-making, but also for reconsidering what exactly the BG learns. Feedback based learning in BG pathways has been best described as an Actor-Critic process (Sutton & Barto, 1998) where the values of alternative actions are learned by trial-and-error comparison of an action’s expected and observed values. The Actor learns to select more valuable actions based on the feedback from the Critic about the difference between expected and observed rewards following an action. Thus, the critical learning signal in RL models is quantified as a reward prediction error (RPE), calculated as the difference between an action’s observed and expected value. Evidence from human and animal studies has consistently linked this form of learning to phasic modulation of dopaminergic neurons in the substantia nigra pars compacta (SNc) that send feedback signals to striatal direct and indirect MSNs. When an action is followed by an unexpected reward (i.e. a positive RPE), SNc neurons display a transient burst in firing that scales with the RPE magnitude, causing proportional influx of dopamine into the striatum. In contrast, the omission of an expected reward (i.e., a negative RPE) cause a transient pause in SNc firing, thereby reducing dopamine availability in the striatum. Recent computational and experimental studies have started to build a more complete picture of the interface between controlled action decisions, as discussed in the previous section, and the role of dopamine in flexibly adapting goal-directed behavior. In the following section we discuss a reconceptualized model of cortico-BG pathways at the intersection of neurocomputational theories of decision-making and reinforcement learning.

### 3.1 Dopaminergic modulation of Believer-Skeptic balance

Electrophysiological studies have consistently found a relationship between the phasic activation of midbrain dopaminergic neurons and the trialwise magnitude of RPEs that mediate action-value RL. For this dopaminergic RPE to be a viable learning signal it must be capable of selectively encouraging rewarded actions and discouraging unrewarded or punished actions. The phasic increase in dopamine following a surprising reward both sensitizes dMSNs and desensitizes iMSNs, making it easier for cortical inputs to quickly execute that action in the future (Hart, Rutledge, Glimcher, & Phillips, 2014; Wiecki & Frank, 2013). By the same token, phasic dips in dopamine following the omission of an expected reward offset the balance in the other direction, requiring stronger or prolonged cortical input to gate the same action in the future (Bahuguna et al., 2015; Gurney et al., 2015; Marcott, Mamaligas, & Ford, 2014). The bidirectional effect of positive and negative feedback on pathway-specific neural subtypes sheds light on the utility of selecting actions with two opposing pathways instead of a single facilitation pathway (Hart et al., 2014). Indeed, several lines of evidence suggest that dopaminergic modulation of the direct pathway is primarily driven by positive RPEs, facilitating approach-learning, whereas the modulation of the indirect primarily is primarily driven by negative RPEs, facilitating avoidance learning (Cox et al., 2015; Frank, Doll, Oas-Terpstra, & Moreno, 2009; Hikida et al., 2013).

In a series of computational experiments, Gurney, Humphries, and Redgrave (2015) recently provided a comprehensive description of the interactions between tonic and phasic fluctuations in striatal dopamine that guide goal-directed action selection. In their neural network model, cortical nput from competing sensory populations is sent in parallel to all three cortico-BG pathways epresenting the sensory-paired actions. Thus, when sensory information is equivocal and cortical input leads to comparable activation in different action channels, the history dependent corticostriatal weights are what critically determine which of the two actions wins out in the selection process.

The synaptic tuning of these weights by positive and negative RPEs can be naturally incorporated into the Believer-Skeptic decision network shown in Figure 2A – by increasing the sensitivity of the direct and indirect populations following rewarded and punished actions, respectively. Over the course of several trials, the feedback-dependent tuning of synaptic weights leads to faster gating in the network and thus faster rates of evidence accumulation in decision space for higher valued actions. This is captured in Figure 4A where the model gradually learns the relative value of alternative actions based on probabilistic stimulus-reward contingencies trial-and-error feedback. Here, similar to the behavioral paradigm used by Frank et al., (2004) the model is presented with a pair of stimuli and must learn to select the stimulus with a higher probability of yielding a reward. Each stimulus is converted into an action by a corresponding pair of direct and indirect nodes that are tuned by corrective feedback signals, simulating the effects of dopaminergic RPE signals on dMSNs and iMSNs. Thus, feedback sensitizes the direct pathway and suppresses the indirect pathway for the optimal choice while shifting the balance in the opposite direction for the alternative, converging on weights that reflect the expected difference in their learned values. In the accumulator model, this manifests as a drift-rate for each stimulus proportional to its perceived value, leading to a stronger choice bias when deciding between alternatives that are less evenly matched in terms of their expected payout (Figure 4A).

**Figure 4.**
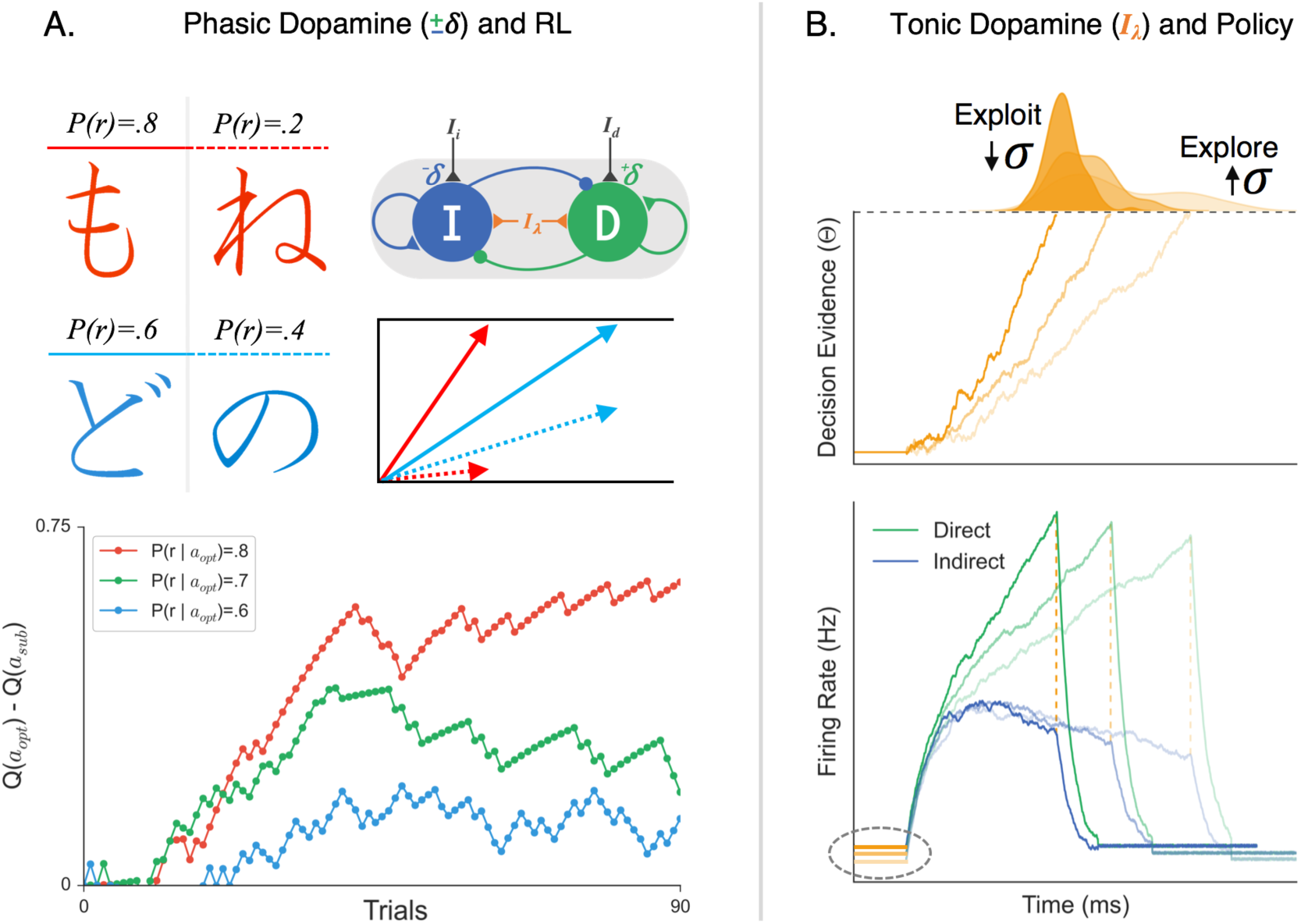
The effects of phasic and tonic dopamine on Believer-Skeptic competition. **(A)** Simulation of probabilistic value-based decision task (upper-left; see Frank, Seeberger, & O’Reilly, 2004) in which the agent must learn the relative value of two arbitrary stimuli based on trial-and-error feedback. On each trial the agent makes a decision by choosing between a pair of Japanese symbols, one with a higher probability of yielding a reward (left column; chosen with action *a*_*opt*_) than the other (right column; chosen with action *a*_*sub*_). Value-based decisions are simulated as a race-to-threshold between two stochastic accumulators (see Figure 3), each reflecting the direct-indirect competition within a single action channel (see Figure 2). Both actions start out with equal associated values Q(*a*_*opt*_)=Q(*a*_*sub*_) and thus, equal drift-rates of accumulation. On each trial, the corrective effects of phasic changes in dopamine are simulated by enhancing (depressing) the sensitivity of the direct (indirect) pathway following positive outcomes (+δ) and vice-versa following negative outcomes (δ). In the accumulator model, this learning results in an increase in the drift-rate for *a*_*opt*_ (solid arrow) and a decrease in the drift-rate for *a*_*sub*_ (dotted arrow), proportional to the difference in their associated value. The bottom panel shows the timeline of the estimated value difference for alternative actions (Q(*a*_*opt*_) - Q(*a*_*sub*_)) for three different probabilistic reward schedules. Stimulus pairs with a greater discrepancy in reward probability (i.e., red > green > blue) lead to faster associative value learning. *(B)* Simulated effects of tonic dopamine levels on exploration-exploitation tradeoff. Tonic dopamine levels were simulated by varying the strength of non-specific background inputs (I_λ_) in a network with stronger weighting of cortical input to direct than indirect pathway. Bottom panel: the same ratio of cortical input to the direct (green) and indirect (blue) pathways leads to faster gating in the presence higher I_λ_ (darker colors, increased baseline) compare to when I_λ_ is low (lighter colors, decreased baseline). Top panel: Increasing tonic levels of I_λ_ facilitates exploitation of the current cortico-striatal weights by accelerating evidence accumulation, resulting in faster decisions and reduced trial-to-trial variability in RT. In contrast, behavior is substantially more variable with lower levels of I_λ_, promoting an exploration policy.

Because in this example the stimulus-action-value associations are probabilistic, a certain amount of exploration is needed in order to optimize the estimated value for each of the two stimuli. In Actor-Critic RL, exploratory decisions are determined by a simple epsilon value that determines the initial probability of going with the currently highest-valued option. Here, however, exploration is naturally handled by the stochastic nature of the direct-indirect competition during the decision process. A recent study found that the RT distributions of value-based choices in a perceptual learning experiment were well described by a DDM in which the learned value difference between alternative stimuli determined the drift-rate of accumulation (Frank et al., 2015). This finding adds support to the future hybridization of RL and decision models, suggesting that the behavioral dynamics of value-based choices can be systematically characterized by corrective modulation of a stochastic rise-to-threshold process.

In addition to the phasic dopamine modulations responsible for learning action-value associations, the level of tonic dopamine availability in the striatum has recently been proposed to regulate the tradeoff between exploratory and exploitative learning policies (Humphries, Khamassi, & Gurney, 2012; Kayser, Mitchell, Weinstein, & Frank, 2015). That is, in order to maximize rewards in dynamic environments (with changing response-outcome contingencies), one must balance the time spent exploring the value of novel, potentially high-payoff actions and exploiting historically rewarding actions (Humphries et al., 2012; Keeler, Pretsell, & Robbins, 2014a). Put into the context of the Believer-Skeptic framework, explorative states can be thought of as conditions in which the balance is tipped towards the Skeptic such that all action possibilities are uncertain and thus no single decision dominates. In contrast, exploitative states are those in which the Believer dominates for a single decision, resulting in faster and more precise decisions that preclude alternative actions from being engaged.

Much of the current understanding of the interplay between value-based learning mechanisms and exploitation/exploration tradeoff policies has come from research on song-bird learning (Brainard & Doupe, 2002; Kao, Doupe, & Brainard, 2005). While research on song-bird learning has progressed largely in parallel with the studies of decision-making in the BG, it has been speculated that the two fields are currently moving towards a mutually beneficial junction (Ding & Perkel, 2014). Juvenile song-birds initially learn to sing by mirroring the song of an experienced tutor but over time compose an individualized version of the song by sampling alternate spectral and temporal components of vocalization (Tumer & Brainard, 2007). This is done to improve reproductive success, as females tend to select males with unique songs that can be performed repeatedly with high precision. Recently, Woolley et al., (2014) found that when practicing in isolation, males express substantially more variability in the spectral and temporal dimensions of song vocalization than when in the presence of a mate. This contextual alternation between exploring alternate song renditions during practice and exploiting a favorite rendition led to systematic differences in the variability of firing in the output of a region called Area X, a homologue of the mammalian BG. The authors proposed that social context led to a changes in the tonic level of dopamine available to neurons in the input structure of Area X, similar to the striatum of the BG, which impacted the amount of exploration or exploitation of the system. Their hypothesis was supported by the observation that striatal connections exhibit a many-to-one convergence onto target cells in the BG output nucleus. Previous work suggests, that given this many-to-one motif, enhanced dopaminergic tone would establish a more consistent average level of activation within a group of striatal units, thus increasing reliability of temporally-locked bursts and pauses of neurons in the output nucleus (Tumer & Brainard, 2007)

Consistent with a dopaminergic regulation between explore-exploit policies, several recent computational modeling studies have found that the simulated effects of tonic dopamine level have a marked impact on action variability (Klanker, Feenstra, & Denys, 2013; Morita & Kato, 2014; Yawata, Yamaguchi, Danjo, Hikida, & Nakanishi, 2012). Increasing dopaminergic availability in the striatum leads to a general “Go” bias in the network, due to the inverse effects of dopamine on MSN subpopulations. Furthermore, higher tonic dopamine levels also increases D1 and D2 receptor occupancy so that RPE signals communicated by phasic bursts and pauses in SNc fail to have the same impact on cortico-striatal plasticity (Keeler, Pretsell, & Robbins, 2014b). Thus, behavior is stabilized to promote exploitation of previously learned associations by facilitating BG throughput that reflects the present weighting scheme at cortico-striatal synapses. In Figure 4B, the population firing rates are shown for different decision policies, all reflecting the same ratio of input to the direct and indirect pathways, but with a change in background levels of tonic dopamine (e.g., background excitation). Increasing dopamine reduces the time constant of evidence accumulation so that learned cortico-striatal weights can be exploited to rapidly accelerate the network toward a “Go” state, with little variability in the RT and outcome of the decision process (Fig. 4B). Alternatively, the same levels of cortical input leads to substantially greater trial-to-trial variability in decision behavior when dopamine is scarce, demonstrated by the widening of the RT distribution for decisions made under lower levels of background dopamine. When considered in the context of selecting from multiple actions, the increase in action variability (i.e., wider RT distribution) with reduced levels of tonic dopamine would allow the agent to explore novel, potentially more rewarding, stimulus-action associations. When a sufficiently rewarding association is found or when there is a change in context that demands precision, increasing background dopamine levels would temporarily halt feedback-dependent plasticity to ensure lower variability in performance.

The relationship between action variability and striatal dopamine adds an interesting perspective to recent studies showing how behavioral variability expands and contracts with a subject’s learning rate, and seems to do so in a controlled, systematic fashion. While standard RL models assume learning rate to be constant, indexing an individual’s inherent sensitivity to feedback error, applying this assumption to human behavior seems to be overly restrictive, especially in realistically dynamic environments. It has been hypothesized, that in settings with a high probability of experiencing a state change (i.e., change in a previously learned stimulus-response-outcome mappings) humans may deliberately amplify the uncertainty or perceived risk of their surroundings so as to maximize adaptability to new information (O’Reilly, 2013). Indeed, a recent study by Wu and colleagues (2014) found that when learning to make visually-guided and reward-guided reaching movements, human subjects demonstrated a simultaneous increase in learning rate with movement variability during times of greater uncertainty. Incredibly, the authors found that the increase in motor variability was not random, but expressed along task-specific dimensions, suggesting that variability is not only capitalized on but is deliberately employed by the nervous system to facilitate adaptation that minimizes relevant sources of uncertainty. In addition to continuous motor control experiments like this one, discrete choice experiments have found that variability in decision-making strategically fluctuates with model-fit learning rates in response to a shift in the statistics of a task or environment (McGuire, Nassar, Gold, & Kable, 2014; Nassar et al., 2012; Payzan-LeNestour, Dunne, Bossaerts, & O’Doherty, 2013).

This idea is supported by a recent fMRI study showing that activity in the caudate nucleus dynamically tracks subjects’ learning rates, rising with greater trial-wise volatility in choice difficulty across blocks of a Stroop task (Jiang, Beck, Heller, & Egner, 2015). The authors found that the volatility-driven changes in caudate activation resulted from descending control signals in the anterior cingulate cortex (ACC), updating the predicted level of control that would be required for the upcoming trial. One intriguing explanation for this finding is that exploratory dynamics in the striatum are modulated by different sources depending on the dimension of exploration. This is driven by changes in background cortical excitation when exploring along the dimension of control and by tonic levels of dopamine when exploring along the dimension of value (Woolley et al., 2014). Of course, this mechanism is only speculative and future studies investigating the mechanisms of control‐ and value-based exploration will need to draw on evidence from both animal models as well as human neuroimaging experiments. Neuroimaging experiments in particular could be poised to investigate this question by comparing the functional connectivity between cortical regions such as ACC and preSMA and the striatum across conditions in which task performance relied on state-change detection of stimulus-control or stimulus-value associations. Furthermore, pharmacological manipulations could be employed to determine if value-based exploration is selectively impaired by increasing tonic dopamine availability.

## 4. Summary & Conclusions

The emerging evidence on the organization of and interactions between BG pathways highlights the limitations of the canonical model of parallel, independent cortico-BG pathways. While the canonical model continues to provide a valuable benchmark for evaluating advancements in the understanding BG function, recent evidence suggests that competition between BG pathways has profound implications for understanding the BG’s role in decision making and learning. Here, we have presented an overview of recent experimental and computational evidence for a reconceptualized view of cortico-BG pathways, highlighting three central themes 1) the direct and indirect pathways engage in competition during action selection, acting as weights on a pulley, rather than independent facilitation and suppression levers, 2) this competition is critical for integrating contextual uncertainty (i.e., Skeptic) with accumulating evidence (i.e., Believer) during decision making and 3) this competitive dynamic lays the foundation for a rich, flexible behavioral repertoire when combined with the dopaminergic modulation described by Actor-Critic RL theories. Based on these findings we have outlined a conceptual framework for the decision-making computations embedded in the competition between the direct and indirect pathways. We feel that this Believer-Skeptic framework offers an appealing first step toward synthesizing neurocomputational theories of decision making with Actor-Critic models of RL.

## Acknowledgments

We would like to thank Julie Fiez and Brent Doiron for their thoughtful feedback regarding the structure and content of the manuscript. This work was funded by the National Science Foundation CAREER Award #1351748.

